# Assembly of von Willebrand Factor Tubules with *in Vivo* Helical Parameters Requires A1 Domain Insertion

**DOI:** 10.1101/2022.07.05.498862

**Authors:** Gabriel Javitt, Deborah Fass

**Affiliations:** Department of Chemical and Structural Biology, Weizmann Institute of Science, Rehovot 7610001 Israel

**Author notes:** Correspondence: Gabriel Javitt or Deborah Fass, Department of Chemical and Structural Biology, Weizmann Institute of Science, Rehovot 7610001 Israel.

## Abstract

The von Willebrand factor (VWF) glycoprotein is stored in tubular form in Weibel-Palade bodies (WPBs) prior to secretion from endothelial cells into the bloodstream. The organization of VWF in the tubules promotes formation of covalently linked VWF polymers and enables orderly secretion without polymer tangling. Recent studies have described the high-resolution structure of helical tubular cores formed *in vitro* by the D1D2 and D′D3 amino-terminal protein segments of VWF. Here we show that formation of tubules with the helical geometry observed for VWF in intracellular WPBs requires also the VWA1 (A1) domain. We reconstituted VWF tubules from segments containing the A1 domain and discovered it to be inserted between helical turns of the tubule, altering helical parameters and explaining the increased robustness of tubule formation when A1 is present. The conclusion from this observation is that the A1 domain has a direct role in VWF assembly, along with its known activity in hemostasis post-secretion.

**Key points:**

- A cryo-EM structure shows that the A1 domain is necessary for forming VWF helical tubules matching those in Weibel-Palade bodies.
- The A1 domain has a role in intracellular VWF supramolecular assembly in addition to platelet binding following secretion and activation.

## Introduction

The polymeric plasma glycoprotein von Willebrand factor (VWF) comprises multiple domains, some of which are responsible for its supramolecular assembly and secretion and some for its function in platelet recruitment and coagulation. The carboxy-terminal region of VWF forms disulfide-bonded dimers as the first step of multimerization.^1,2^ The amino-terminal region, which consists of three VWD (D) assemblies (Figure 1A), subsequently links these dimers together to form polymers.^1,3^ The middle region, containing three tandem VWA (A) domains, carries out activities required for hemostasis. The VWA1 (A1) domain has binding sites for platelet glycoprotein Ibα (GPIbα), heparin, and collagens.^4–7^ A1 has also been proposed to bind growth factors and thereby promote wound healing.^8^ The VWA2 (A2) domain has an ADAMTS13 protease cleavage site to aid in clearance of VWF from the body and can undergo force-induced unfolding.^9,10^ The VWA3 (A3) domain contains a collagen-binding site that contributes to initial VWF tethering at sites of vascular injury.^11^

**Figure 1.**
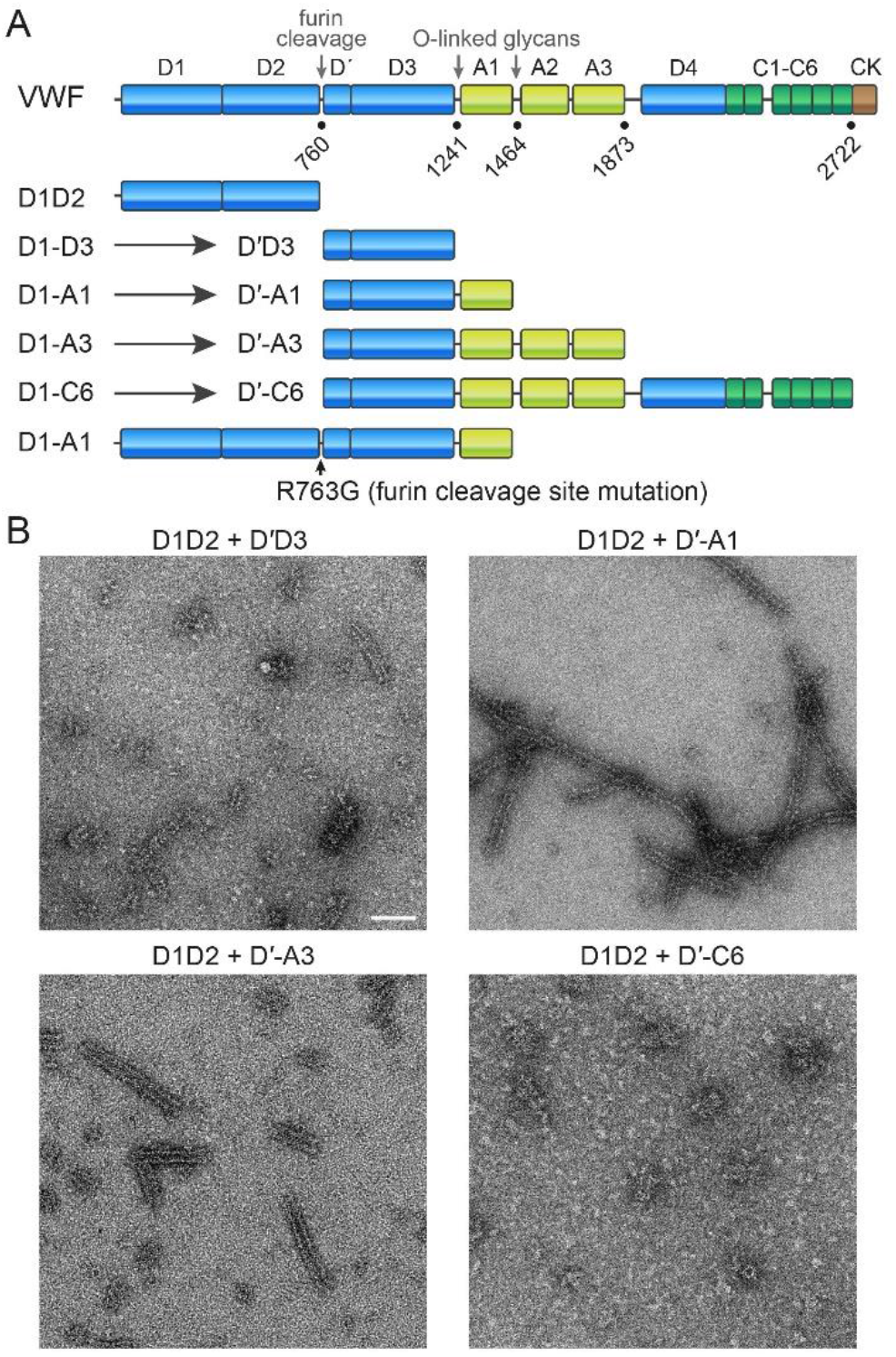
VWF domain map and segments produced for supramolecular assembly. (A) The primary structure of VWF is displayed with the major assemblies/domains labeled. Amino acid residue numbers at boundaries of segments studied herein are indicated. Also shown are the VWF segments produced and analyzed in this study. All expression plasmids encoded segments beginning with D1, but, due to furin cleavage (except in the R763G mutant), protein fragments beginning at D′ were obtained for segments extending past D2. (B) Electron micrographs of negatively stained tubular assemblies formed upon incubation of the indicated VWF segments. Scale bar is 100 nm and applies to all micrographs.

A striking feature of VWF is its propensity to form tubules, which are stored in intracellular compartments of endothelial cells called Weibel-Palade bodies (WPBs), prior to VWF secretion into the bloodstream. The amino-terminal portion of VWF, reconstituted from the D1D2 and D′D3 fragments (Figure 1A), spontaneously forms tubules *in vitro* in an acidic pH range characteristic of the Golgi apparatus and WPBs.^12^ The structure of these tubules has been determined, first at low resolution^12^ and then at higher resolution using cryo-electron microscopy (cryo-EM).^13,14^ In addition, the organization of VWF in intracellular WPBs was studied by cryo-EM tomography and Fourier analysis.^15^ In all these cases, and in structural studies of filaments formed by the amino-terminal region of the intestinal mucin, MUC2,^16^ which is homologous to VWF, the fundamental unit of the supramolecular assembly was a globular “bead.” MUC2 filaments are made of linear strings of these beads,^16^ whereas the VWF tubule is made of successive beads spiraling around a central axis.^12^ These beads-on-a-string or beaded tubule arrangements promote disulfide bond formation between amino-terminal domains during the construction of disulfide-bonded polymers.^16^ After secretion, the beads disassemble to yield head-to-head and tail-to-tail disulfide-linked linear polymers.^17^

It was previously observed that the presence of the A1 domain in recombinant VWF amino-terminal fragments led to more robust tubule formation *in vitro*^12^ and that the A1 domain was required for tubule formation *in vivo*.^18^ We sought to determine the structural basis for this observation. One possible explanation would be that A1 packs against the outside of the D-assembly tubule, stabilizing its structure. We found, however, that A1 directly inserts into the tubule, explaining the discrepancy between the geometries of *in vitro*^12–14^ *vs. in vivo*^15^ VWF tubules.

## Methods

### Protein production and purification

The coding sequence for VWF D1D2 (UniProt ID P04275 residues 1 to 745) was inserted into the pCDNA3.1 vector with a carboxy-terminal His_6_ tag as described previously.^13^ For other constructs, the full-length VWF gene was amplified from clone 100068741 of hORFeome5.1 with primers containing an amino-terminal AflII site and a carboxy-terminal His_6_ tag coding sequence plus an EcoRI site and inserted into pCDNA3.1. D1-D3 (residues 1 to 1241 of the VWF preprotein), D1-A1 (residues 1 to 1464), D1-A3 (residues 1 to 1873), D1-C6 (residues 1 to 2722) were generated by deletion of the unwanted DNA from the parental plasmid using the Q5 Site-Directed Mutagenesis Kit (NEB BioLabs). The R763G point mutation was introduced into D1-A1 by PCR mutagenesis. Plasmids were propagated in *Escherichia coli* XL-1 cells and verified by sequencing.

Proteins were produced by transient transfection of plasmids into HEK 293F cells using the polyethylenimine (PEI) Max reagent (Polysciences, Inc.) with a 1:3 ratio (wt/wt) of DNA to PEI at a concentration of 1 million cells per ml. Cells were maintained in FreeStyle 293 medium. Six days after transfection, the culture medium was collected and centrifuged for 10 min at 500 g to pellet cells. The supernatant was then centrifuged for 20 min at 4,000 g to pellet any remaining particulate matter. The supernatant from this second centrifugation was filtered through a 0.45-μm filter, and the His_6_-tagged proteins were purified by nickel-nitrilotriacetic acid (Ni-NTA) chromatography. Proteins were concentrated to 0.6 mg/mL and dialyzed against a buffered solution containing 20 mM 4-morpholineethanesulfonic acid (MES), pH 5.6, 10 mM CaCl_2_, and 150 mM NaCl. Expression of D1-D3, D1-A1, and D1-C6 led to purification of D′D3, D′-A1, and D′-C6, respectively, due to furin cleavage between D2 and D′ (Figure 1A). The carboxy-terminal fragments were retained during Ni-NTA purification as they contained the His_6_ tag.

### Sample preparation for electron microscopy

For negative staining, purified VWF proteins including A1 domains were dialyzed to 0.3 mg/mL in 50 mM MES, pH 5.9, 150 mM NaCl, 10 mM CaCl_2_ at a 2:1 molar ratio of VWF D1D2 to VWF D′-A1 and incubated at 37 °C for 24 hr. VWF proteins containing more domains did not form tubules under these conditions, but did form tubules in 50 mM MES, pH 5.9, and 10 mM CaCl_2_. After incubation, proteins were diluted in their respective buffers to 0.03 mg/mL, and 3 μl was applied to a glow discharged (PELCO easiGlow) carbon-coated 300 mesh copper grid (Electron Microscopy Sciences). Grids were stained with 2% uranyl acetate and allowed to dry. Samples were visualized using a Tecnai T12 electron microscope (Thermo Fisher Scientific) equipped with a OneView camera (Gatan). For cryo-EM, 3 μl of each incubated protein solution was applied to glow discharged UltrAuFoil R 1.2/1.3, 300 mesh gold grid (Quantifoil). Using a Vitrobot Mark IV plunger (Thermo Fisher Scientific), grids were plunge frozen from a chamber held at 10 °C and 100% humidity into liquid ethane cooled by liquid nitrogen.

### Cryo-EM image acquisition

Cryo-EM data were collected on a Titan Krios G3i transmission electron microscope (Thermo Fisher Scientific) operated at 300 kV. Movies were recorded on a K3 direct detector (Gatan) installed behind a BioQuantum energy filter (Gatan) using a slit of 20 eV. Movies were recorded in counting mode at a nominal magnification of 105,000×, corresponding to a physical pixel size of 0.83 Å. The dose rate was set to 19 e^−^/pixel/sec, and the total exposure time was 1.5 sec (accumulated dose ~41 e^−^/Å^2^). Each movie was split into 40 frames of 0.0375 sec. Nominal defocus range was −0.5 to −2 μm. EPU (Thermofisher) was used for automated data collection, in which a single image was collected from the center of each hole. Image shift was used to navigate within 10-μm arrays and stage shift to move between arrays.

### Cryo-EM image processing

Recorded movies were imported into CryoSPARC v3.3.1 and subjected to Patch-Based motion correction and Patch-Based contrast transfer function (CTF) estimation.^19^ Micrographs with better than a 4.0-Å CTF fit resolution were retained for further processing. For VWF D1D2+D′-A1 this resulted in 3,387 micrographs retained from an initial 4,368. For VWF D1D2+D′-A3 6,064 micrographs were retained from 6,756. For VWF D1-A1 R763G 3,778 were retained from 5,169. Particles were picked using either the template picker or the filament tracer with 30-40 Å separation distance between boxes. Initial numbers of overlapping particles were 739,073 for D1D2+D′-A1, 10,292,183 for D1D2+D′-A3, and 311,132 for D1-A1 R763G. Extensive two-dimensional classification yielded 457,730 particles for D1D2+D′-A1, 63,772 particles for D1D2+D′-A3, and 101,153 particles for D1-A1 R763G. These particles were used for *ab initio* helical refinement performed with constraints on in-plane rotation and longitudinal shifts but no additional symmetry, starting with an initial helical twist of 83° and helical rise of 28.7 Å. D1 symmetry was then imposed, and helical parameters were further refined. Particles were symmetry expanded with the resulting helical parameters, and local refinement was performed without symmetry using a mask encompassing two successive beads along the helical path or two beads between helical turns. A 20° limit was placed on the rotation search in the local refinement to prevent overfitting.

### Model building and structure refinement

To generate a model for the tubules, a two-bead reconstruction for human VWF D1D2 +D’D3 (PDB ID 7PMV) was rigid body fit into the local refinement map of two successive beads along the helical path using the UCSF Chimera tool ‘fit in map’.^20^ The module was refined into the density using Coot full chain refinement.^21^ The A1 domains (PDB ID 7A6O) were rigid body fit into the local refinement map of two beads between helical turns. The helical refinement and local maps were combined into one focused map with Chimera ‘vop maximum’ for refinement. The initial models were rebuilt and refined by iterative cycles of Phenix real-space refinement,^22^ interactive rebuilding in Coot, and ISOLDE,^23^ and validated using Molprobity.^24^ Model refinement statistics are summarized in supplemental Table 1. UCSF ChimeraX^25^ and Pymol^26^ were used for figure generation. Interface areas were calculated using Pymol. The cryo-EM density maps and atomic coordinates of D1D2+D′-A1 tubules have been deposited in the Electron Microscopy and Protein Data Banks with accession codes EMD-14998 and PDB 7ZWH. The cryo-EM density maps for D1D2+D′-A3 and D1-A1 R763G have been deposited in the Electron Microscopy Data Bank with accession codes EMD-15004 and EMD-15005, respectively.

## Results

VWF tubules were reconstituted *in vitro* by incubating the D1D2 propeptide^27^ together with the D′-A1 segment, the D′-A3 segment, or the D′-C6 segment (Figure 1A) at pH 5.9. Tubules were also produced from the D1-A1 segment containing the R763G mutation, which disrupts the furin cleavage site and leaves the propeptide attached (Figure 1A).^28^ The resulting tubules were compared to those generated in previous studies using the D1D2 and D′D3 segments.^13,14^ The tubules were longer when the A1 domain was present, as reported also by others,^12^ but were shorter with the inclusion of the A2A3 segment, and shorter still with the inclusion of the C domains, as seen by transmission electron microscopy (TEM) of negatively stained samples. (Figure 1B).

Cryo-EM data were collected on tubules containing the A1 domain and those containing all three A domains. Sections of the tubules were selected from the images, these image regions were classified, and three-dimensional reconstructions were performed (Figure 2A). In the maps generated for tubules containing the D′-A3 segment, density was seen for the A1 domain but not for the A2 or A3 domains (not shown). This observation indicates that A2 and A3 do not participate in tubule formation. In all maps, the resolution of the A1 domain was lower than the core of the D assemblies but similar to other surface-exposed regions (Figure 2A).

**Figure 2.**
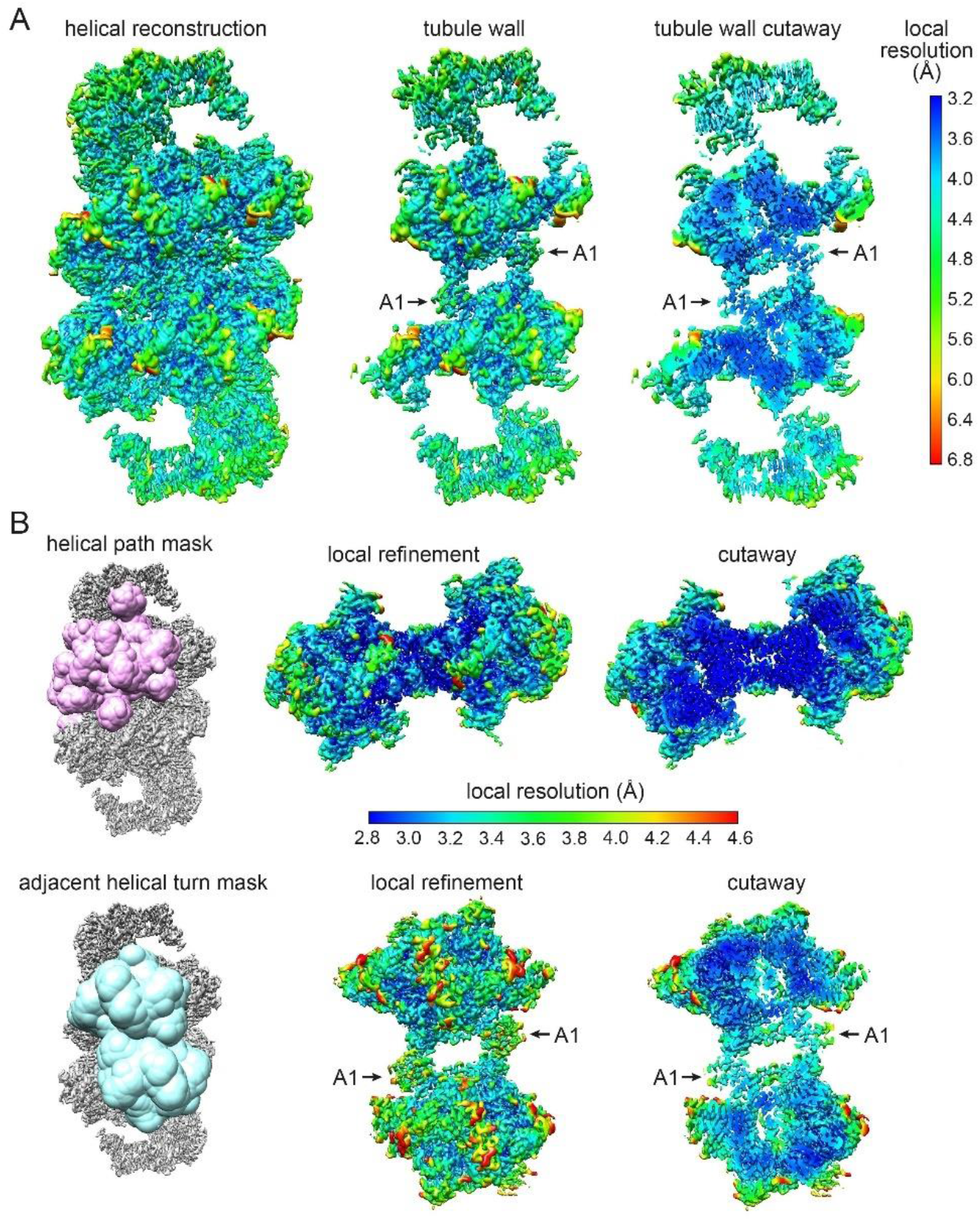
VWF tubule maps. (A) VWF map obtained by helical reconstruction, colored according to local resolution. The left panel shows the full map, the central panel is a slab containing the front wall of the tubule, and the right panel is a cutaway view of the tubule wall. (B) Maps obtained by local refinement of the indicated masked regions. The resolution scale bar applies to the local refinement maps for both masks.

From the reconstruction of tubules formed using the D′-A1 segment, masks encompassing bead pairs either along the helical path or between helical turns were designed, and local refinement was performed to achieve higher resolution for contacts between neighboring beads (Figure 2B). Reconstructing two adjacent beads along the helical path enabled construction of a model for the entire tubule using the values for rotation and translation between the two beads. Local refinement of a bead pair spanning successive helical turns improved the visualization of the contacts between the A1 domains and the tubule beads. In the three-dimensional maps obtained from the two-bead reconstructions, we observed the same internal bead organization seen previously,^13^ with additional density for the A1 domain (Figure 2). Atomic models were built into the maps with the aid of our previous tubule structure composed only of D assemblies^13^ and the known structure of the A1 domain (*e.g*., PDB ID codes 7A6O, 1SQ0, 1UEX, 1IJK).

Overall, the tubules with and without the A1 domain appeared quite similar (Figure 3A). However, the A1 domain did not merely dock onto the D-assembly tubule but rather replaced the interhelical contacts observed previously.^13,14^ In the structure of the D-assembly tubule, these contacts occurred by reciprocal interaction of the C8-1/TIL1 regions.^13^ In the tubules containing A1, the A1 domain was found wedged between the E1 module in one bead and the C8-1 region of another bead (Figure 3B). Notably, with the A1 domain present, it was possible to obtain a map at a nominal resolution of 3.2 Å (supplemental Figure 1) for a pair of beads interacting between helical turns, whereas previous local refinements of such bead pairs yielded maps to only about 4.5 Å resolution.^13,14^

**Figure 3.**
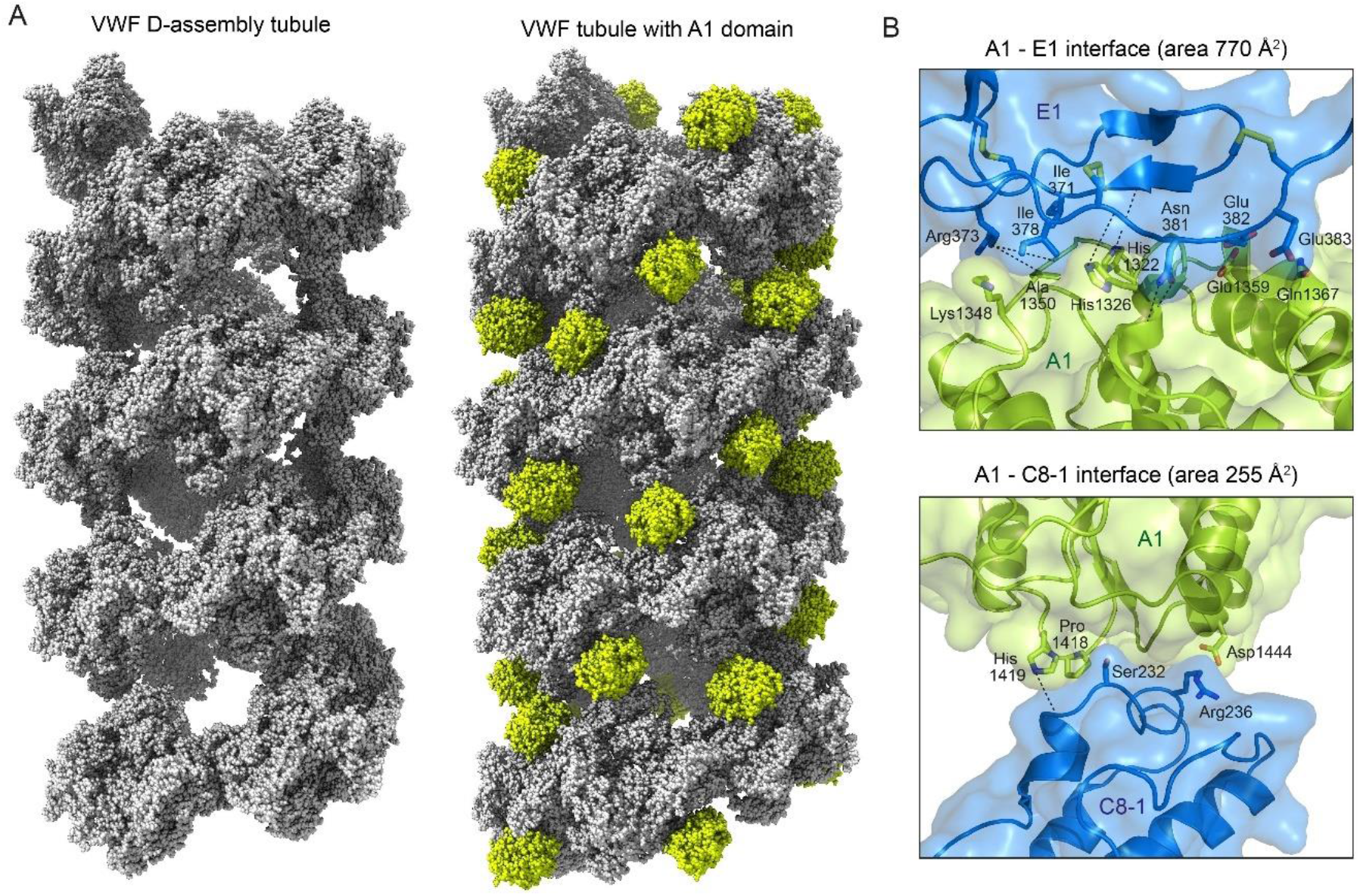
The A1 domain in VWF tubules. (A) Atomic sphere representation of D-assembly tubules (left)^13^ and tubules containing the A1 domain, colored chartreuse (right). (B) Details of the two interfaces between A1 and the D-assemblies in the tubule. Dashed lines represent potential hydrogen bonds between side chains and backbone atoms, which are not shown explicitly in the ribbon diagram.

The presence of the A1 domain within the tubule explains the difference in helical parameters between the *in vitro* tubules generated from D assemblies and the full-length tubules studied in cells (Table 1). Fourier analysis of the tubules in WPBs showed an axial rise per subunit of 27.7 Å.^15^ The axial rise per subunit of the *in vitro* D-assembly helical tubules was 28.5 Å,^13^ whereas the axial rise of the in *vitro* tubules containing A1 described here was 27.6 Å, consistent with the *in vivo* tubules (Table 1). While it may seem counterintuitive that insertion of an extra domain shortens the axial rise, the explanation is that the helix unwinds slightly in the presence of A1 (supplemental Movie 1). The helical twist (*i.e*., rotation angle around the helix axis) between beads was 86.0° for the D-assembly tubule and 83.1° for the tubule with the A1 domain. Overall, the tubules lacking and containing the A1 domain have similar pitches: 119 Å for the D-assembly tubule and 120 Å for the tubule with the A1 domain (Table 1).

**Table 1:** Helical parameters for VWF tubules.

| VWF variant | axial rise per subunit (Å) | helical twist (°) | pitch (Å) | reference |
| --- | --- | --- | --- | --- |
| in cells | 27.7 | 83.1 | 120 | ref. 15 |
| D1D2 - D'D3 | 26.2 | 85.6 | 110 | ref. 12* |
| D1D2 - D'D3 | 28.5 | 86.0 | 119 | ref. 13 |
| D1D2 - D'D3 | 29.4 | 86.8 | 122 | ref. 14 |
| D1D2 - D'D3A1 | 27.6 | 83.1 | 120 | this work |
| D1D2 - D'D3A1A2A3 | 27.6 | 82.6 | 120 | this work |
| D1D2D'D3A1 R763G | 28.1 | 82.6 | 122 | this work |

Highlighted rows emphasize the close agreement between the high-resolution VWF tubule structure reported in this work and the parameters measured from full-length VWF tubules in WPBs in endothelial cells.
\* Heterogeneity in pitch was observed in this study. Values in this table correspond to the major bin used for reconstructing the VWF low-resolution structure.

The map resolution at bead interfaces was sufficient to appreciate specific interactions that stabilize the tubule, both between beads along the helical path and longitudinally *via* the A1 domain. The interactions previously observed between neighboring beads in the spiral of the VWF D-assembly tubule^13^ were maintained in the tubule containing the A1 domain. Specifically, cation-π interactions described between the inter-bead bridge and the D1 and D2 assemblies within a bead (Y730 and R202, F716 and H460, and F722 and R442) were unaffected by the present of the A1 domain. In contrast, the small interfaces between the C8-1 and TIL1 regions in the D-assembly tubules^13^ were disrupted by insertion of A1. A new interface was formed between the A1 domain and the E1 module, rather than the TIL1 module, of the D1 assembly. This new interaction is related to the unwinding of the helical tubule in the presence of A1 (supplemental Movie 1).

The A1-E1 interface is mostly charged and polar, consistent with pH-dependent assembly, though some hydrophobic contacts are also seen, including the packing of the aliphatic part of the Lys1348 side chain against Ile378 and the Cα of Ala1350 against Ile371. The charged and polar interactions include contacts between His1322 and His1326 in A1 with backbone carbonyls in E1. In addition, Gln1367 in A1 interacts with Glu382 in E1, and Glu1359 is near Glu383, potentially making carboxyl-carboxylate interactions at low pH.^29^ Arg373 in E1 contacts backbone carbonyls in A1. Finally, Asn381 forms hydrogen bonds with an edge strand of the β-sheet, strand β3, in A1. The area of the interface between A1 and E1 is 770 Å^2^ (buried surface area 1539 Å^2^).

The second interface, made between A1 and the C8-1 region, is smaller, with an interface area of 255 Å^2^ (buried surface area 509 Å^2^). This interface contains an interaction between His1419 in A1 and a backbone carbonyl in C8-1, and a potential salt bridge between Asp1444 and Arg236. Hydrophobic contacts are made between Pro1418 in A1 and the Cα and Cβ of Ser232 in C8-1, as well as the Cβ of Ala235. As beads interacting longitudinally along the tubule axis are bridged by two A1 domains, the total interface area between A1 domains and bead pairs is 2048 Å^2^ (buried surface area 4098 Å^2^). These values are almost four times those measured for directly interacting beads from successive turns of the D-assembly VWF helical tubule, which had an interface area of 520 Å^2^ (buried surface area 1040 Å^2^).^13^

Many of the interface residues described above for the tubule containing A1 are more highly conserved in VWF than His288, which was present in the interbead interface along the tubule axis in the D-assembly tubule.^13^ For example, Glu1359 is almost invariant in VWF, and Arg373 is very highly conserved (occasionally replaced by lysine or glutamine). Interface residues Arg236, Lys1348, Glu382, and His1322, are also well conserved and, when substituted, are replaced by amino acids with similar properties. Though A1 amino acids may also be conserved for downstream functional reasons, the conservation of key amino acids participating in A1 interfaces in the tubule and the congruence of helical parameters with tubules in WPBs indicate that the A1-containing tubules are an accurate representation of the physiological VWF tubule core.

Despite the overall high resolution of the maps and clear density for most portions of the protein in the tubules, including the A1 domain, density was missing for certain segments. Specifically, as previously observed for D-assembly tubules,^13^ density was absent in all maps for the region of the furin cleavage site (residues 741-766), even in the tubules prepared from protein in which the site was mutated to prevent cleavage. Density was also absent for the region between D3 and the A1 domain (residues 1198-1264) (Figure 4A). In the tubular arrangement, two beads in successive helical turns interact *via* two A1 domains (Figure 2). Residue 1197, at the end of the modeled region of D3, was at comparable distances from the two A1 domains, such that the D3-A1 connectivity was ambiguous (Figure 4B). In contrast to a ~40 amino-acid linker between the D3 assembly and the following domain in the mucin MUC2, which appeared as contiguous density in 2D classes and the 3D reconstruction of cryo-EM data,^16^ the VWF tubule maps did not reveal the direction of the D3-A1 linker. However, the dimensions of the missing E3 module of the D3 assembly, together with about 23 amino acids likely to assume extended conformations due to high proline content and O-glycosylated threonines,^30^ would be sufficient to span the gap to either A1 domain.^31^

**Figure 4.**
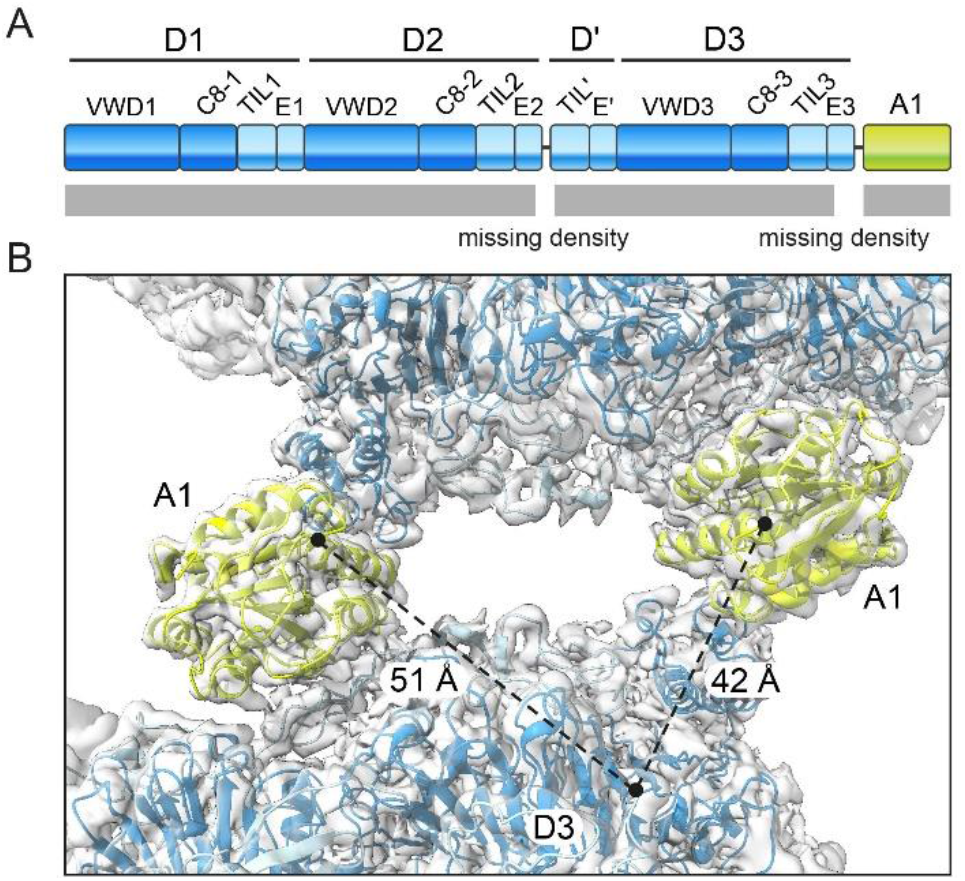
Disordered linker between D3 and A1. (A) Map of the domain structure of the VWF amino-terminal segment showing the disordered regions in the tubule (missing density). (B) The endpoint of a D3 assembly modeled into the tubule map (residue 1197) and the startpoint of the two nearest A1 domains (residue 1263) are indicated by black circles. Dashed lines indicate the two possible connectivities with distances indicated. The shorter distance is not necessarily the more likely to represent a covalent connection.

Structures are available of the A1 domain in complex with various binding partners, allowing comparison between the interfaces used to incorporate A1 into the tubule and those used for downstream function. Particularly interesting is a comparison with the binding mode of A1 to GPIbα. The structure of a complex between A1 and GPIbα (PDB code 1SQ0)^32^ revealed that the A1 β-sheet edge-strand β3 makes four hydrogen bonds in an intermolecular β-sheet formed with the GPIbα switch region (Figure 5). Though this mode of interaction is different from the interaction of A1 with the rest of the tubule (Figure 3B), and the E1 and GPIbα footprints on A1 are not identical (Figure 5), A1 uses the β3 edge strand both for contacts within the tubule and for adhesion to platelet surface receptors. Therefore, participation of A1 in the tubule hides its GPIbα binding surface.

**Figure 5.**
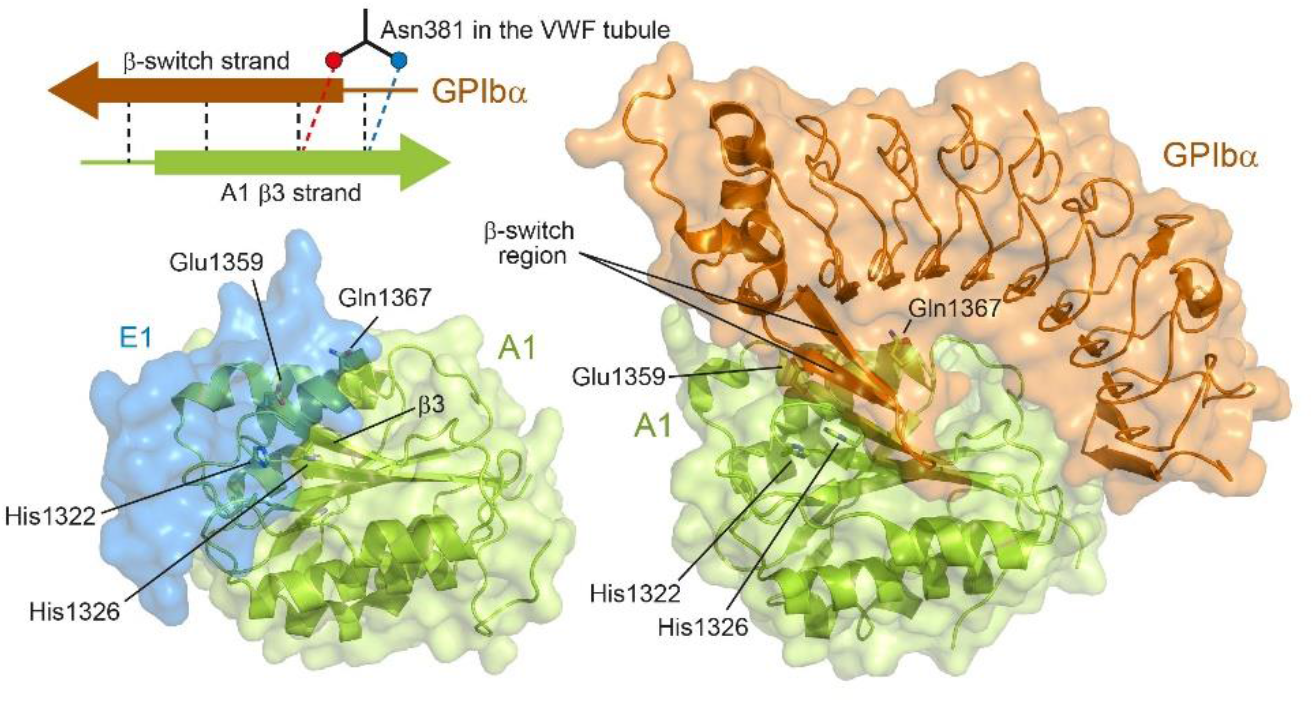
Comparison of A1 tubule binding and GPIbα binding. In the upper left is a diagram of hydrogen bonding interactions (dashed lines; not to scale) made by the A1 edge strand with either Asn381 in the VWF tubule or with a β-switch strand in the GPIbα complex. The two hydrogen bonds made with the Asn381 side chain are replaced by inter-strand hydrogen bonds in the GPIbα complex. In the structure images, A1 is shown looking down the central β-sheet. Left, a semitransparent surface of E1 (blue) is shown to indicate where it contacts A1. Right, the complex between A1 and GPIbα (PDB code 1SQ0).

Additional A1 residues involved in the interaction with GPIbα are also buried in the VWF tubule. Adjacent to the E1 residue Asn381, which caps the A1 β3 strand in the tubule, is Glu382, which contacts Glu1359 in A1. This Glu-Glu contact is likely to require the low pH of the Golgi and WPBs. In the complex with GPIbα, Glu1359 instead contacts a tyrosine hydroxyl, an interaction consistent with a neutral pH environment. Gln1367, which contacts Glu383 in the tubule, interacts with threonine and phenylalanine side chains in the GPIbα complex. Disruption of nearby clusters of acidic residues may contribute to breaking the interaction between Asn381 and the A1 edge strand upon secretion of VWF into the neutral pH environment of the bloodstream. However, as the process of secretion does not activate A1, other interactions are expected to protect the interface until A1 is triggered for GPIbα binding.

## Discussion

The A1 domain is critical for VWF function. A1 binds collagen exposed at wound sites and GPIbα on platelets, helping to initiate platelet plug formation and activate the clotting cascade^33,34^. Many von Willebrand disease mutations map to the A1 domain.^35^ These mutations either compromise or increase platelet binding, or they influence the amount of high molecular weight multimers found in the circulation through effects on VWF production or turnover.^35^ The importance of A1 to VWF function is reflected in the intensity of research efforts to uncover A1 activation mechanisms and to control A1 binding activity to produce antithrombotic therapies.^36–40^

The major finding presented here is that, prior to secretion, the A1 domain is integrated into the VWF helical tubule core. The significance of this discovery is two-fold. One, it explains the observation that tubules formed by VWF segments containing the A1 domain are both more robust^12^ (Figure 1B) and slightly different from^15^ those composed of D assemblies only.^13,14^ Two, it suggests a mechanism for intracellular shielding of key A1 binding surfaces. A major functional region of A1 is the edge-strand β3, which is central to the interaction of VWF with GPIbα (Figure 5).^32^ We found that β3 is hidden in the tubule and that its exposed edge is protected by hydrogen bonding to a well-conserved amino acid in the D1 assembly (Figure 5). Other conserved amino acids, including the site of the von Willebrand disease mutation Glu1359Lys in A1,^41^ also participate in shielding the GPIbα binding site of A1 within the tubule.

The observation that A1 is integral to the VWF tubule core complements a study of the VWF A1-CTCK segment (Figure 1A) and an analysis of the “dimeric bouquet” formed by this region.^31^ The glycosylated segments flanking A1 were proposed to provide the flexibility necessary to flatten the A1-CTCK bouquet against the helical tubule core, as required for close lateral packing of tubules in WPBs.^15^ The knowledge that A1 is immobilized within the tubule, and the invisibility of A2 and A3 in reconstructions of tubules made with the D′-A3 segment, indicates that the required flexibility comes from the glycosylated segment carboxy-terminal to A1. The glycosylated segment amino-terminal to A1 may be useful for other purposes, including enabling A1 to assume the position and orientation necessary to insert into the tubule.

A major focus of current research is on the role of A1 flanking regions in modulating A1 properties.^37,38,42-44^ For example, the flanking regions have been proposed to interact cooperatively to shield the A1 GPIbα binding site.^37^ These segments were not detected in the cryo-EM maps of our tubules, even when both the amino- and carboxy-terminal flanking segments were present, as in the D1D2+D′-A3 tubules. The interactions proposed between the segments flanking A1 may be too dynamic to detect by cryo-EM. Alternatively, they may form after tubule disassembly upon secretion into the bloodstream, where conditions differ from the low-pH incubations used to mimic the VWF assembly and storage environments in the Golgi apparatus and WPBs. In general, release of A1 from its site in the tubule may be required for interaction with other segments of the VWF glycoprotein. The tubule structure containing A1 serves as an accurate starting point for addressing the question of whether and how A1 interactions in the tubule are replaced by binding with, or shielding by, other parts of the glycoprotein to generate the latent form of VWF in the circulation.

The findings presented herein demonstrate definitively that the A1 domain participates in formation of VWF tubules that match the properties of the full-length protein in WPBs inside endothelial cells.^15^ The residue-resolution A1-containing tubule structure (Figure 3A, right) can thus be considered a model of the physiological VWF tubule core. The observation made by multiple groups^12–14^ that VWF segments lacking the A1 domain nevertheless form helical tubules *in vitro* with certain similarities (*i.e*., diameter, pitch, start symmetry) to *in vivo* VWF tubules^15^ reinforces the notion we put forth previously: filaments with the symmetry of VWF and mucins^16^ lend themselves to higher-order organization into helices due to the cooperation of multiple weak interactions along the length of the helical tubule.^13^ In contrast to the small surface areas involving poorly conserved amino acids seen between helical turns in the D-assembly tubules,^13,14^ the A1-containing tubules reported here have larger and better-conserved interfaces along the tubule axis. The key interface involving the A1 edge strand has properties suggesting dependence on low pH, explaining how this substantial interface could be consistent with tubule disassembly upon secretion. In summary, we show that the VWF A1 domain is directly involved in VWF tubule assembly, and likely disassembly, complementing the well-known role of the A1 domain in hemostasis following secretion.

## Acknowledgments

This study was supported by the Israel Science Foundation (grant 2660/20 to D.F.), the Center for Scientific Excellence at the Weizmann Institute of Science (to D.F.), and the Dr. Bruce A. Pearlman Grant for Student-Initiated Research (to G.J.).

## Authorship

Contribution: G.J. produced proteins, determined structures, and analyzed results. D.F. analyzed results. D.F. and G.J. prepared the manuscript.

