## Supplemental data for "Assembly of von Willebrand Factor Tubules with *in Vivo* Helical Parameters Requires A1 Domain Insertion"

### Supplemental Table 1: Data and model statistics

| | |
|---|---|
| Model | VWF D1D2+D´-A1 (PDB ID 7ZWH) |
| Chains | 6 |
| Atoms | 40,421 (Hydrogens: 19,691) |
| Amino acid residues | 2660 |
| Ligands | BMA: 2 |
| | NAG: 12 |
| | Ca: 6 |
| Bonds (RMSD) | |
| Length (Å) (# > 4σ) | 0.003 (0) |
| Angles (°) (# > 4σ) | 0.557 (1) |
| MolProbity score | 2.28 |
| Clash score | 12.38 |
| Ramachandran plot (%) | |
| Outliers | 0.08 |
| Allowed | 15.62 |
| Favored | 84.30 |
| Rotamer outliers (%) | 0.26 |
| Cβ outliers (%) | 0.00 |
| Peptide plane (%) | |
| *Cis* proline/general | 2.9/0.0 |
| Twisted proline/general | 0.0/0.0 |
| CaBLAM outliers (%) | 7.95 |
| B-factors (min/max/mean) | |
| Protein | 48.90/304.40/126.47 |
| Ligand | 96.13/261.07/155.21 |
| Data | |
| Box | |
| Lengths (Å) | 176.02, 127.68, 172.30 |
| Angles (°) | 90.00, 90.00, 90.00 |
| Model vs. Data | |
| CC (mask) | 0.81 |
| CC (box) | 0.67 |
| CC (peaks) | 0.38 |
| CC (volume) | 0.81 |
| Mean CC for ligands | 0.83 |

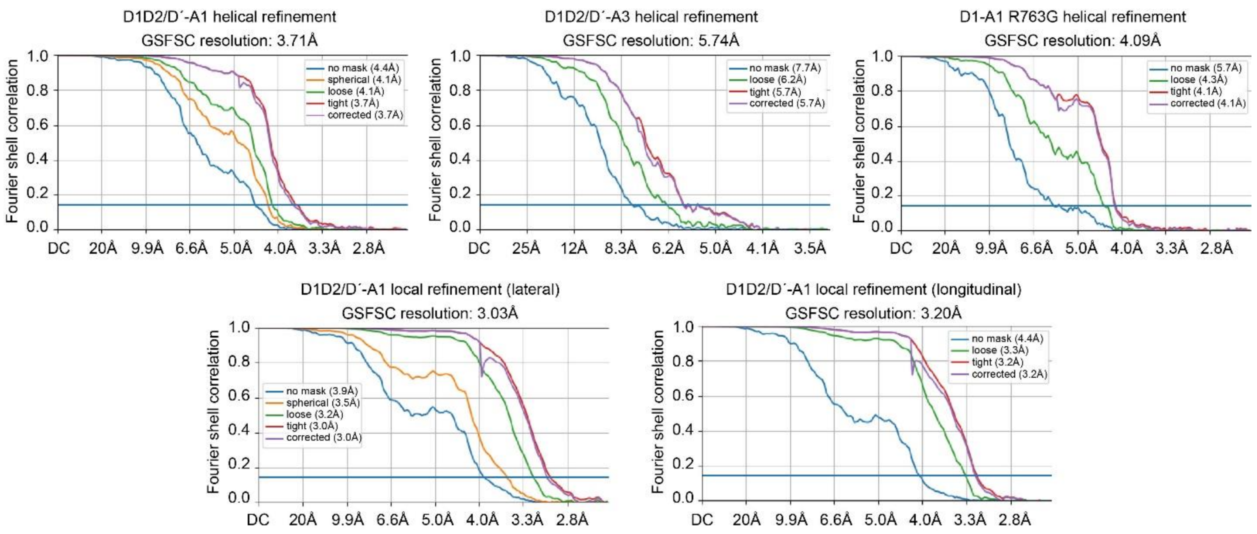


**Supplemental Figure 1. Fourier shell correlation (FSC) curves.** Resolution at a FSC cut-off of 0.143 is shown above each plot. DC is direct current, the zero-frequency component of the Fourier transform.

**Supplemental Movie 1. Conformational differences between VWF tubules with and without A1.** A morph animation shows the conformational differences between tubules constructed from VWF D assemblies alone (PDB ID 7PNF) and tubules containing the A1 domain (colored chartreuse). The apparent unwinding of the helix in the presence of A1 is due to the participation of the E1 module, rather than the TIL1 module, in interturn contacts in the helix.
